# Transcriptome analysis reveals higher levels of mobile element-associated abnormal gene transcripts in temporal lobe epilepsy patients

**DOI:** 10.1101/2021.05.14.444199

**Authors:** Kai Hu, Ping Liang

## Abstract

**Objective:** To determine role of abnormal splice variants associated with mobile elements in epilepsy.

**Methods:** Publicly available human RNA-seq-based transcriptome data for laser-captured dentate granule cells of post-mortem hippocampal tissues from temporal lobe epilepsy patients with (TLE, N=14 for 7 subjects) and without hippocampal sclerosis (TLE-HS, N=8 for 5 subjects) and healthy individuals (N=51), surgically resected bulk neocortex tissues from TLE patients (TLE-NC, N=17). For each individual sample, *de novo* transcriptome assembly was performed followed by identification of spliced gene transcripts containing mobile element (ME) sequences (ME-transcripts) to compare the ME-transcript frequency across the sample groups. Enrichment analysis for genes associated with ME-transcripts and detailed sequence examination for representative epileptic genes were performed to analyze the pattern and mechanism of ME-transcripts on gene function.

**Results:** We observed significantly higher levels of ME-transcripts in the hippocampal tissues of epileptic patients, particularly in TLE-HS. Among ME classes, SINEs were shown to be the most frequent contributor to ME-transcripts followed by LINEs and DNA transposons. These ME sequences almost in all cases represent older MEs normally located in the intron sequences, leading abnormal splicing variants. For protein coding genes, ME sequences were mostly found in the 3’-UTR regions, with a significant portion also in the coding sequences (CDS) leading to reading frame disruption. Genes associated with ME-transcripts showed enrichment for involvement in the mRNA splicing process in all sample groups, with bias towards neural and epilepsy-associated genes in the epileptic transcriptomes.

**Significance:** Our data suggest that abnormal splicing involving MEs, leading to loss of function in critical genes, plays a role in epilepsy, particularly in TLE-HS, providing a novel insight on the molecular mechanisms underlying epileptogenesis.

**Key points box:**

- Significantly higher rates of abnormal splicing variants involving mobile elements (MEs) were observed in the hippocampal tissues of epilepsy patients.
- SINEs/Alus are most frequently observed in ME-transcripts followed by LINEs and DNA transposons.
- For protein coding genes, MEs mostly locate in 3’ UTR, but also in coding regions, causing open reading frame disruption, with a bias for neural and epileptic genes.
- Abnormal splicing involving MEs may be a contributing factor in epileptogenesis.

## 1. Introduction

Epilepsy is one of the most common and serious neurological diseases with mesial temporal lobe epilepsy (MTLE) as the most common form of epilepsy, showing a frequent pathological feature of hippocampal sclerosis and for being the most conventional forms of drug-resistant epilepsy group ^1^. Although the disease-causing factors for epilepsy are diverse and heterogeneous, epilepsy is widely considered as a highly genetic and heritable condition under many situations ^2–4^. In a review by Wang et al.,^5^ a total of 977 genes was compiled as epilepsy genes, neurodevelopment-associated genes, epilepsy-related genes or potential epilepsy-associated genes. Epilepsy genetics presently focuses on protein-coding genes, more specifically concentrating on ion channel and neural receptor mutations, which mostly lead to idiopathic epilepsies, as well as gene copy number variations (CNVs) led by deletions and duplications that cause epilepsy and brain developmental disorders ^6, 7^. This has led to a few limitations of current epilepsy genetics studies. For example, clinical genetic analysis is limited to variants near or within genes with a known association in human epilepsy disease with a focus on those in the coding regions, leading to the use less than 1% of the genome sequencing data, such as those from next generation sequencing, for clinical diagnosis ^8, 9^. As a result, the expression and function of non-coding sequence, as a major component of the genome being largely ignored and unexplored in prior epilepsy genetic studies. Mobile elements (MEs), also known as transposable elements (TEs), constitute at least 50% of the human genome ^10–12^. MEs are categorized as DNA transposons and retrotransposons with the latter further divided into long terminal repeats (LTRs), long interspersed nuclear element (LINE) class of autonomous retroelements, short interspersed nuclear element (SINE) ^13^, and SINE-VNTR-Alus (SVA) ^14^ with SVA being a very young family unique to the hominid primate group ^15^. MEs participate in gene function and regulation related to many biological processes, such as neurogenesis, brain development, and aging via a plethora of mechanisms including interruption of protein coding, alteration of RNA splicing or gene regulation ^16–18^. Through insertional mutation at the germline level, MEs are reported to be responsible for ~1% of human genetic diseases ^19, 20^, including a spectrum of central nervous system (CNS) diseases, such as amyotrophic lateral sclerosis (ALS), Alzheimer’s disease (AD), Parkinson’s disease (PD), schizophrenia, bipolar disorder, and post-traumatic stress disorder (PTSD) ^21–23^. Furthermore, emerging evidence also suggests a possible role of somatic retrotransposons by LINE/L1s in brain development and in neuronal diseases ^24^. For epilepsy, limited studies have also implied a role of MEs via ME-mediated genomic rearrangements in CDKL5 and ALDH7A1^25, 26^ and *de novo* somatic L1 insertions biased towards to epilepsy associated genes ^27^. However, a general landscape depicting the involvement of MEs in epilepsy is still missing.

As an attempt to better understand the molecular mechanism involving MEs in contributing to epilepsy, in this study we examined the involvement of MEs in RNA splicing and showed a higher presence of MEs in gene transcripts leading to abnormal splicing in the hippocampal tissues of TLE patients, particularly in TLE-HS, than the normal control and control brain tissues. Our study represents the first study suggesting the ME-mediated abnormal RNA splicing as a contributing factor in epileptogenesis.

## 2. Materials and methods

### 2.1. Sources of RNA sequences for human epilepsy patients

For this study, we used existing RNA-seq data for laser-captured dentate granule cells of post-mortem hippocampal tissues from the mesial temporal lobe epilepsy (MTLE) patients with and without hippocampal sclerosis (HS) generated by Griffin et al. ^28^ (NCBI BioProject accession: PRJNA290212). The subject number for MTLE without hippocampal sclerosis (TLE) and with hippocampal sclerosis (TLE-HS) is 7 and 5, respectively, with the sample size being 14 and 8, respectively, due to replicated samples taken from some subjects. We also included the RNA-seq samples of bulk neocortex tissues from MTLE patients (TLE-NC) as a type of negative tissue control at a sample size of 17 from the datasets of Kjær et al. ^29^ (NCBI BioProject accession: PRJNA556159). Another RNA-seq dataset for laser-captured dentate granule cells of post-mortem hippocampal tissues from healthy individuals was chosen as the normal control at a sample size of 51 (NCBI BioProject accession: PRJNA600414). Based on the original descriptions, all laser captured hippocampal tissues were originated from dentate gyrus. The lists of samples used in this study are summarized in Table 1 with the detailed NCBI sample and sequence accessions listed in Table S1.

**Table 1.** Human samples and groups used in this study

| Group Descriptions | Sample description | StudyID (SRA acc#) | #subject | #sample |
| --- | --- | --- | --- | --- |
| Normal control | Laser captured dentate granule cells of post-mortem hippocampal tissues from healthy individuals | PRJNA600414 | 51 | 51 |
| TLE-NC (temporal lobe epilepsy neocortex) | Bulk neocortical tissues from mesial temporal lobe epilepsy patients | PRJNA556159 (Kjær et al., 2019) | 17 | 17 |
| TLE-HS (temporal lobe epilepsy with hippocampal sclerosis) | dentate granule cells of hippocampal tissues from mesial temporal lobe epilepsy patients with hippocampal sclerosis | PRJNA290212 (Griffin NG et al., 2016) | 5 | 8 |
| TLE (temporal lobe epilepsy without hippocampal sclerosis) | Laser captured dentate granule cells of hippocampal tissues from mesial temporal lobe epilepsy patients without hippocampal sclerosis |  | 7 | 14 |

### 2.2. De novo assembly of transcriptome sequences, identification of MEs in the assembled transcripts, and comparison among groups

The RNA-seq data for the samples listed in Table 1 and Table S1 were downloaded from NCBI SRA database (https://www.ncbi.nlm.nih.gov/sra) to Compute Canada high performance computing server (https://computecanada.ca) for *de novo* transcriptome assembly, identification of ME sequences, and position correlation with annotated genes. For protein coding genes, the distribution of MEs in transcripts is also further broken down into 5’ UTR, CDS, and 3’ UTR regions. The number of gene transcripts containing ME sequences (ME-transcripts) in each sample was normalized as the number of ME-transcripts per million transcripts (TPM). TPM values were also calculated for each major ME class (SINE, LINE, LTR, SVA, and DNA transposons) and for their subfamilies. More related details are provided in Supplemental Methods, while the statistical analysis was performed as described in section 2.5.

### 2.3. Functional enrichment analysis of genes associated with ME-transcripts

To find out whether the presence of the ME-transcripts show any bias towards certain types of genes, we performed enrichment analysis for genes associated with ME-transcripts for each sample group individually using the DAVID tool (https://david.ncifcrf.gov) ^30^. Specifically, a non-redundant list of genes associated with ME-transcripts in all individual samples for a sample group was generated and used for enrichment analysis. For enrichment analysis, gene ontology (GO terms), KEGG pathways, and additional functional categories including UP_KEYWORDS, UP_SEQ_FEATURE, AND UP_TISSUES provided by the DAVID tool were included. We collected all categories showing statistically significant enrichment with a Benjamini adjusted p-value below 0.05.

### 2.4. Identification of epilepsy-associated genes and neural genes associated with ME-transcripts

We collected a list of 977 genes known to be associated with epilepsy compiled by Wang et al. ^5^ and a list of 2,449 genes associated with neurogenesis and neural system development from the MGI database (http://www.informatics.jax.org). The numbers of epilepsy and neural genes associated with the ME-transcripts were collected for each sample by cross-mapping between the genes associated with the ME-transcripts and the epilepsy associated and neural gene lists for comparison among groups. For a selected number of ME-transcripts associated with epilepsy genes, we performed detailed sequence analysis to predict the functional impact of the ME sequence on the host genes. For the *SCN1A* gene, the expression level of the ME-transcript in comparison with the normal splicing forms was analyzed and compared across sample groups. More detailed method descriptions for this part is provided in the Supplementary Methods.

### 2.5. Statistical analysis

Data analysis and figure plotting were performed by using a combination of R package, GraphPad Prism, and Microsoft Excel. Statistics were performed using SAS 9.4 (SAS Inc, Cary, NY, USA). Data were reported using descriptive statistics. Variables were presented as arithmetic mean plus standard deviation (mean±STD). To assess the statistical significance of the variables across different sample groups, one-way analysis of variance (ANOVA) followed by Student-Newman-Keuls post hoc tests (SNK-q test) were used for those showing normal distribution, while Krustal-Wallis rank sum test followed by Nemenyi test or Wilcoxon rank sum test and were used for those showing none-normal distribution, and Chi-square test was used for categorical variables. All p-values were from 2-sided tests and results were deemed statistically significant at different levels at p<0.05(**\***), p<0.01(**\*\***), or p<0.001(**\*\*\***).

## 3. Results

### 3.1. Higher rates of ME-transcripts were observed in the hippocampal tissue transcriptomes of human epileptic groups than that of the normal control groups and human TLE neocortex group

As shown in Table 2 and Fig. 1, four classes of MEs, including SINE, LINE, LTRs, and DNA transposon, were observed in the gene transcripts from all sample groups. It is interesting to notice that SVA, as the youngest ME class in the human genome, is completely missing from the ME-transcripts. As shown in Fig. 1A to D, except for SINEs, all other three classes of MEs showed a gradient of the ME frequency from low to high in the order of the normal control, TLE-NC, TLE, and TLE-HS. For SINEs, the frequency in TLE-NC is higher than that of the TLE. However, for all ME classes, it is always that the TLE-HS group has the highest frequency of ME-transcripts, while the normal control group has the lowest, when all transcripts (including non-coding transcripts) were considered, and the differences are statistically significant (p<0.05) for the TLE and TLE-HS groups in comparison with the normal control group for all four ME classes (Fig. 1A to D). It is worth noting that except for SINEs, the TLE-NC group, which was included as a control brain tissue from the epilepsy patients but irrelevant to epilepsy, has a ME-transcript frequency that is lower than that of the TLE and TLE-HS, and being higher but without significant difference from the normal control (Fig. 1B-D).

**Table 2.** Contribution of mobile elements (MEs) to gene transcripts in human epilepsy groups

| Genic region<br>ME class | All exon regions |  |  |  | CDS |  |  |  |
| --- | --- | --- | --- | --- | --- | --- | --- | --- |
|  | control | TLE-NC | TLE | TLE-HS | control | TLE-NC | TLE | TLE-HS |
| SINEs | 3734.8 ± 963.1 | 7664.4 ± 4374.6 | 6466.5 ± 3476.4 | 9983.4 ± 3734.5 | 196.0 ± 70.6 | 189.1 ± 165.6 | 242.6 ± 181.9 | 347.1 ± 272.5 |
| LINEs | 1916.9 ± 715.4 | 2774.2 ± 2036.5 | 4397.8 ± 2009.1 | 5117.5 ± 2294.6 | 173.5 ± 114.8 | 61.8 ± 61.5 | 139.3 ± 83.4 | 140.5 ± 99.2 |
| LTRs | 746.0 ± 255.2 | 979.2 ± 653.8 | 1513.5 ± 927.0 | 1804.5 ± 836.0 | 37.2 ± 24.9 | 109.0 ± 25.0 | 54.0 ± 39.7 | 25.3 ± 22.0 |
| DNA transposons | 1270.1 ± 529.3 | 2103.7 ± 1324.8 | 2560.4 ± 1166.3 | 3186.9 ± 1619.5 | 69.1 ± 40.2 | 69.2 ± 45.4 | 68.4 ± 56.2 | 108.3 ± 76.1 |
| All MEs | 7751.3 ± 2249.5 | 13615.9 ± 8156.9 | 15113.6 ± 6891.2 | 20277.6 ± 8117.5 | 479.1 ± 175.2 | 317.7 ± 200.5 | 512.1 ± 289.8 | 598.8 ± 399.4 |

| Genic region<br>ME class | 3' UTR |  |  |  | 5' UTR |  |  |  |
| --- | --- | --- | --- | --- | --- | --- | --- | --- |
|  | control | TLE-NC | TLE | TLE-HS | control | TLE-NC | TLE | TLE-HS |
| SINEs | 1455.3 ± 439.4 | 3327.1 ± 1994.6 | 2642.3 ± 1416.3 | 4193.4 ± 1552.7 | 51.2 ± 39.6 | 77.6 ± 67.1 | 35.3 ± 33.9 | 46.1 ± 29.2 |
| LINEs | 624.8 ± 240.8 | 1137.6 ± 920.4 | 1761.2 ± 901.7 | 2121.4 ± 1008.0 | 32.8 ± 22.1 | 41.7 ± 35.7 | 50.2 ± 16.0 | 30.0 ± 26.8 |
| LTRs | 237.5 ± 115.1 | 318.1 ± 233.0 | 572.0 ± 399.8 | 741.8 ± 363.7 | 23.7 ± 19.1 | 129.5 ± 66.4 | 13.7 ± 7.1 | 17.3 ± 11.2 |
| DNA transposons | 504.6 ± 226.0 | 929.4 ± 626.9 | 1074.7 ± 502.4 | 1332.3 ± 709.2 | 15.7 ± 13.4 | 39.1 ± 20.2 | 22.2 ± 8.4 | 36.2 ± 32.0 |
| All MEs | 2856.7 ± 908.5 | 5755.9 ± 3644.2 | 6128.5 ± 2887.1 | 8474.5 ± 3412.4 | 115.7 ± 62.5 | 137.9 ± 104.3 | 47.7 ± 42.5 | 91.1 ± 73.8 |

The frequency of MEs was calculated as the average number of MEs per million assembled transcripts within the sample group with the standard deviation also provided. Values on bold print are those showing statistically significant differences between control and the epileptic groups.

**Figure 1.**
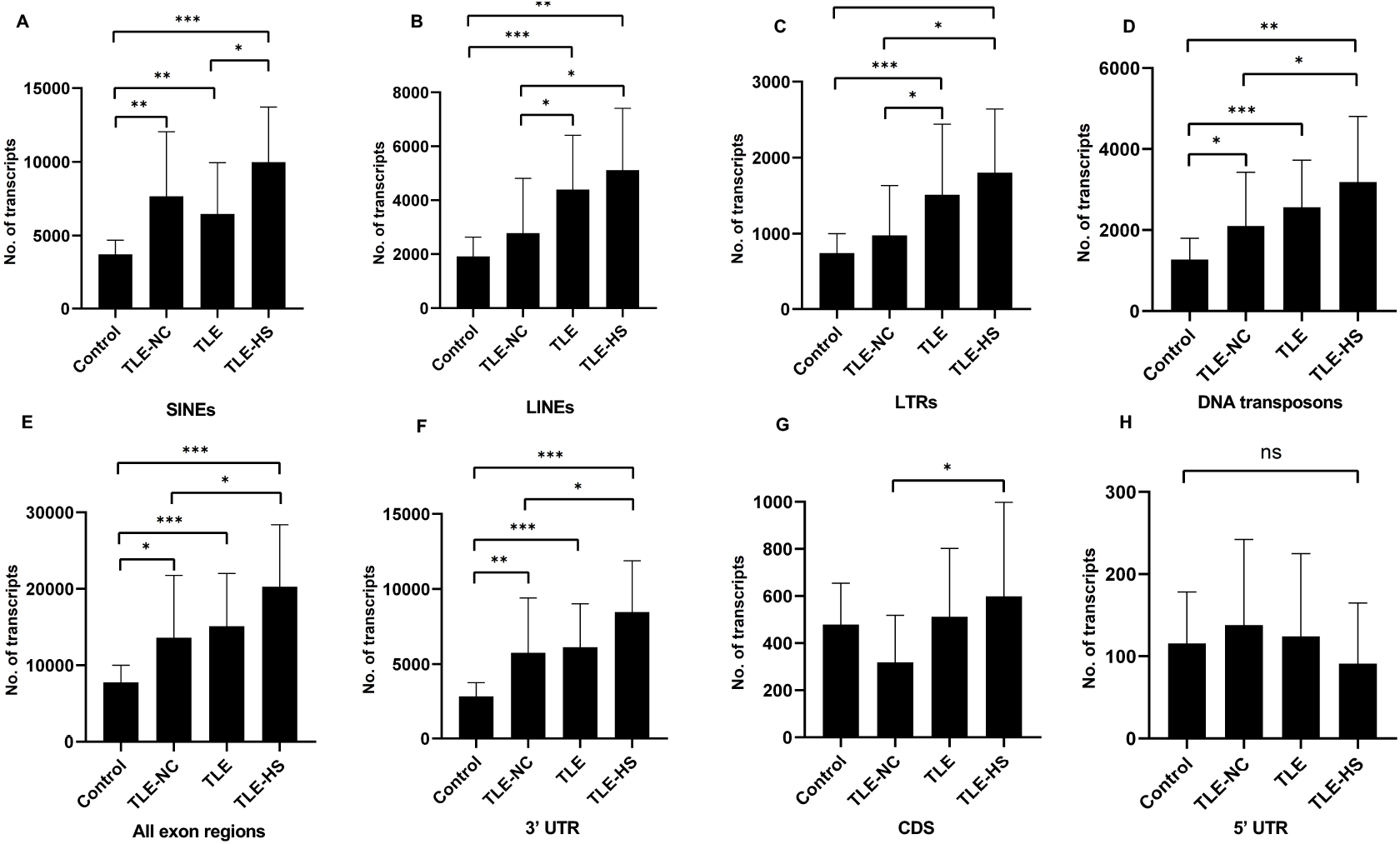
Differential expression of MEs in gene transcripts among different human epileptic groups. The frequency of MEs in gene transcripts (all exon regions) in each sample group is expressed as the average number of MEs per million assembled transcripts for SINEs (A), LINEs (B), LTRs (C), and DNA transposons (D) in different human epileptic groups. The frequency of MEs in gene transcripts (all exon regions) in each sample group is expressed as the average number of MEs per million assembled transcripts in all transcript regions (E), 3’ UTR regions (F), CDS regions (G), 5’ UTR regions (H). The statistical significance of the differences between two sample groups is indicated by the bracket lines at the top of the bar plots with p-value <0.001 indicated by “***”, p-value <0.01 indicated by “**”, p-value <0.05 indicated by “*” above the connecting brackets

We also examined the distribution of MEs across the 3 sub-exon regions of protein coding genes. As shown in Table 2 and Fig. 1E-H, the 3’ UTR region has the highest frequency of ME-transcripts, being more than 10 times higher than that in CDS and 5’ UTR regions. The differential pattern among the sample groups in 3’ UTR region (Fig. 1F) is similar to that of all exon regions (for all genes) (Fig. 1E). For the CDS regions (Fig. 1G), the TLE-HS group has the highest ME-transcript frequency when all ME classes are considered, being significantly higher than that of TLE-NC with the latter being the lowest among the four groups. The 5’ UTR regions had the lowest frequency of MEs among the 3 exon regions, likely due to their much shorter sequences. In contrast to the other two regions and all regions, the ME-frequency in 5’ UTR regions was shown to be the lowest in the TLE-HS group, but with no statistical significance between any sample groups (Fig. 1H).

More detailed comparisons were made among the three exon regions with MEs broken down to ME classes (Table 2). Overall, the frequency of SINEs was always shown to be highest in the entire transcripts and for CDS and 3’ UTR regions. In comparison, the frequency of LINEs was shown to be the highest in TLE-HS only for 3’ UTR. For LTRs, the highest frequency was seen in 3’ UTR of the TLE-HS, but for CDS, its frequency in TLE-HS is the lowest among all sample groups. For DNA transposons, the highest frequency was seen in TLE-HS for all three exon regions (Table 2).

### 3.2. SINEs are the main contributors to ME-transcripts

Among ME classes, SINE is the most frequent in all sample groups, ranging from 3,734 TPM in the control group as the lowest to 9,983 TPM in the TLE-HS group as the highest, when all exon regions were considered (Table 2 and Fig. S1A). This is followed by LINE, DNA, and LTR in order from high to low with the rates of LTRs being more than four times lower than that of SINEs. This order of abundance among ME classes is the same for all sample groups and in all three exon regions of protein coding genes, except for the TLE-NC group, which showed a slightly higher rate of LTRs than DNAs in 5’ UTR and CDS regions (Table 2). The higher rates for SINEs and LINEs can be explained by their higher copy numbers in the human genome (1,779,233 and 1,516,226, respectively ^31^). However, DNA transposons have a lower copy number in the genome than that of LTRs (483,994 vs 720,177 ^31^), and they yet showed a higher rate in ME-transcripts than LTRs (Table 2 and Fig. S1).

To assess and compare the degree of involvement in ME-transcripts proportionally by their abundance in the genome, we normalized the TPM values in Table 2 as TMP per million of MEs (TPMPM) in that ME class in the entire human genome. As shown in Table S2 and Fig. S1A, for all exon regions, after this normalization, while the relative portion dropped, SINE was still shown to be the most frequent ME class, ranging from 3,162.2 TPMPM in the normal group as the lowest to 8,452.8 TPMPM in the TLE-HS group. The relative portion of LINEs and LTRs didn’t seem to change much by normalization (Fig. S1). However, interestingly, the frequency of DNA transposons in TPMPM increased in all sample groups from their TPM values and became the second highest among the ME classes, being higher than that of LINEs and LTRs in all sample groups and very close that of SINEs in the TLE group (Table S2 and Fig. S1A). This indicates that, proportionally, DNA transposons were shown to be active contributors to ME-transcripts than LINEs and LTRs with the largest increase seen in TLE-HS compared to the control group (Table S2 and Fig. S1A). We also examined the situation in the CDS regions and observed a similar increase for DNA transposons when normalized as TPMPM (Fig. S1B), and certainly this would be true also for 3’ UTR regions (data not shown).

We further analyzed the frequency of ME subfamilies within each class and identified subfamilies contributing to differential MEs subfamilies. Among the 36 ME subfamilies detected in epileptic transcriptomes, 10 subfamilies showed significant difference in their frequencies in ME-transcript at least for one pairwise group comparison. As shown in Table S3, the 10 subfamilies include Alu, MIR from SINEs, L1, L2, and CR1 from LINEs, ERVL-MaLR, ERV1, and ERVK for LTRs, and hAT-Charlie, TcMar-Tigger, and hAT-Tip100 for DNA transposons in order from high to low within their respective ME classes.

By age of MEs, the overall pattern seems to be that MEs involved in ME-transcripts are limited to members from older subfamilies. This is also in agreement with the absence of SVAs as the youngest ME class in these ME-transcripts. However, a higher portion of younger members was observed in TLE-HS than the control for LINEs and LTRs (more detailed description is provided in the Supplemental Results and Fig. S2).

### 3.4. Genes associated with ME-transcripts showed enrichment for involvement in RNA splicing in epileptic groups and for epilepsy-associated and neural genes

To find out whether the impact of MEs on RNA splicing is completely random or selective for a certain type of genes, we performed enrichment analysis of genes associated with ME-transcripts for gene function categories including GO terms, KEGG pathways and other category terms provided by the DAVID tool ^30^. The list of categories showing statistically significant enrichment in at least one sample groups were collected and shown in Fig. 2A. It appears that a common theme among the enriched function categories is the mRNA splicing process. Among all sample groups, a list of 206 genes were identified as associated with different components of mRNA splicing based on GO terms, KEGG pathways, and sequence features assigned by the DAVID tool (Fig. 2A). In addition to the first 4 categories with descriptions directly associated with RNA splicing, all remaining categories share most of their genes with these categories. This makes the RNA-splicing process as the single enriched function category, which is very unusual and interesting, and yet very relevant in a sense that these genes were shown to be involved in abnormal splicing. The enrichment for genes expressed in brain tissues was seen in all sample groups, as one can expect, since these genes were detected to be expressed in brain tissues as a starting point of the analysis. Interestingly, among the human sample groups, the TLE-NC (neocortex) group showed the least enrichment for these function categories with none being statistically significant, indicating that neural cells in neocortex are quite different from the those in the hippocampal tissue at least for mRNA splicing.

**Figure 2.**
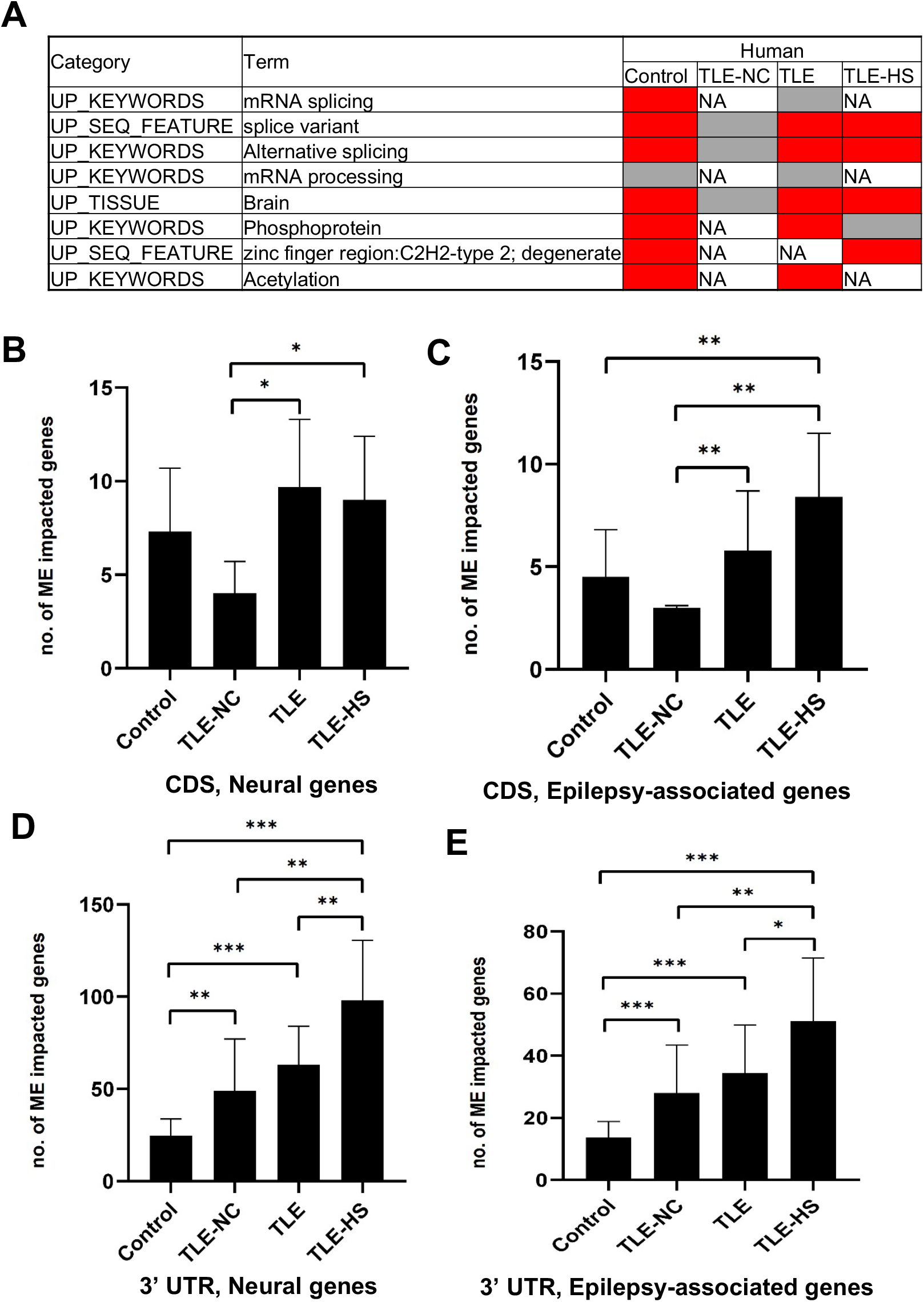
The frequency of mobile elements (MEs) in the transcripts of neural and epilepsy-associated genes and function categories enriched by genes associated with ME-transcripts. A, Function categories enriched by genes associated with ME-transcripts in human and mouse sample groups. Cells in red background indicate enrichment is statistically significant with raw p-value and Benjamini adjusted p-value below 0.05; Cells in grey background indicate enrichment without statistical significance; NA: no enrichment for the category. B-E: Bar plots showing the average of number of neural genes (B) and epilepsy-associated genes (C) in CDS regions and also in 3’ UTR regions (D and E). The statistical significance of the differences between two sample groups is indicated by the bracket lines at the top of the bar plots with p-value <0.001 indicated by “***”, p-value <0.01 indicated by “**”, p-value <0.05 indicated by “*”

We further compared genes associated with ME-transcripts against the lists of epilepsy-associated genes and neural genes to see whether there are any differences between the control and epileptic groups by frequency. Specifically, we calculated the number of genes from each of the two gene lists (2,449 and 977 for neural and epilepsy-associated genes, respectively) in each sample in a group and compared among the groups. In this case, we only included genes with MEs in the 3’ UTR and CDS regions of the transcripts. A non-redundant list of genes associated with ME-transcripts in the CDS regions for the epileptic groups is provided in Table S4.

As shown in Table S5 and Fig. 2C-E, for both neural and epilepsy-associated genes, the TLE-HS group always showed a higher number of genes impacted by MEs than all other groups in both the 3’ UTR and CDS regions. Statistically significant differences were observed for neural genes in CDS regions and for both neural and epilepsy-associated genes in the 3’ UTR regions (Fig. 2B, D, E). It is interesting to note that the TLE-NC group showed the lowest number of genes impacted by MEs in CDS regions from both gene lists with statistical significance from other sample groups (Fig. 2B & C).

### 3.5. MEs involved in abnormal splicing and led to gene function loss

To understand the functional impact of the MEs in gene transcripts, we took genes known to be associated with epilepsy as examples and examined at the detailed sequence level of the transcript. In almost all cases we examined, the MEs in the transcripts represent those normally located in the intron regions but were retained to generate extended exons (exon extension) as shown in Fig. 3 for the *SCN1A* gene. The outcome of such abnormal splicing involving MEs in the CDS region is the disruption of normal open reading frame via generation of a premature stop codon (Fig. 3D). The predicted functional impact for *SCN1A* is a truncated protein product, which misses more than 1000 amino acids at the C-terminus, and/or mRNA degradation by the nonsense-mediated mRNA decay (NMD) machinery, both expected to result in loss of function for the gene.

**Figure 3.**
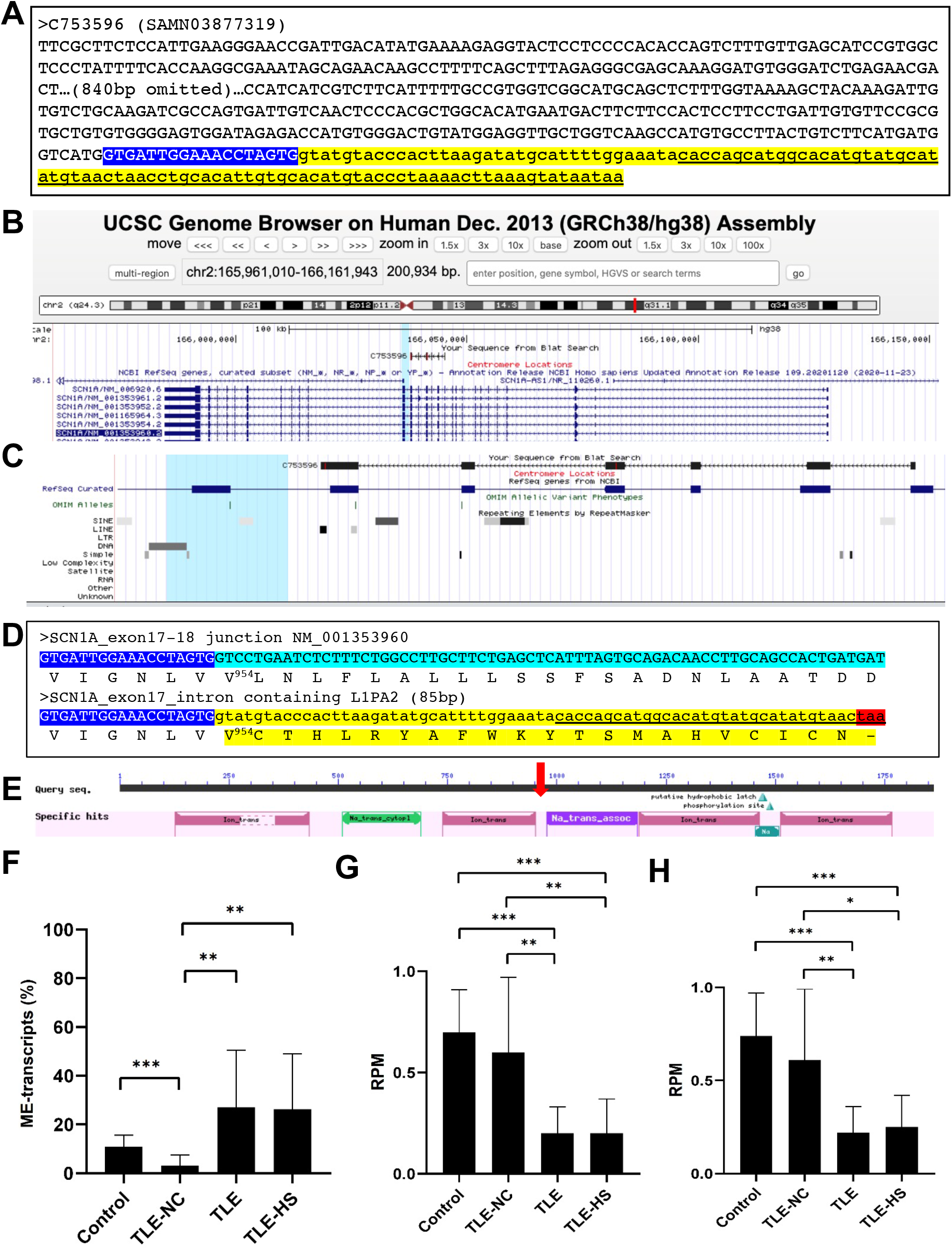
Disruption of open reading frame (ORF) by a ME in the transcript of *SCN1A* gene. **A**. The sequence of a transcript scaffold (C753596 from SAMN03877319) for the *SCN1A* gene from TLE-group in fasta format. The ME sequence is indicated in underlined font in the yellow highlighted part, while the normal exon sequence is labeled in UPPER and the intron sequence in lower case; **B.** A screenshot showing the alignment of the ME-transcript sequence to the human genome in the UCSC Genome Browser, indicating the location of the alignment within the gene structure of the *SCN1A*; **C**. A zoomed-in picture of the alignment of the transcript sequence with the *SCN1A* gene with the Repeat Track at the bottom, showing the ME-transcript sequence covering exon 13 to exon 18 of the gene and the last part of the ME-transcript sequence extending beyond the normal splicing junction of the exon 17 to cover a 87-bp L1PA2 element from the LINE family; **D**. Detailed sequence comparison between the normal transcript (top entry) and ME-transcript (bottom entry) at the junction of the exon 17 & 18 of NM_001353960 with the cDNA sequences shown in the plus strand of the mRNA. The end of the exon 17 sequence is shown in blue background and white font and the first part of the exon 18 sequence shown in green background (top entry), while the intron containing the ME sequence is shown in yellow background (bottom entry). The corresponding protein sequence is provided below the cDNA sequence in each case with the ME-transcript sequence having an early stop codon as indicated by a “-”. The number in superscript indicates the position in the normal protein sequence (NP_001340889, 1980 amino acids in total length) at the point, after which the ME version diverged from the normal version. The colour highlights correspond to those in A; **E**. A screenshot of the NCBI CDD entry for the *SCN1A* protein (NP_001340889) showing the conserved domain structure of the protein with the vertical red arrow indicating the approximate position of the truncation caused by the ME sequence; **F-H,** the expression level of the normal and abnormal ME-transcript for the *SCN1A* gene in the sample groups for the ratio ration between the normal and ME-transcripts (F), the reads per million total reads (RPM) of the normal transcripts (G) and total (H), all based on the counting of raw reads supporting each of the two junction sequences representing the two splice forms. Information on the statistical significance of pairwise comparisons among the sample groups is provided at the top of the plots with p-value <0.001 indicated by “***”, p-value <0.01 indicated by “**”, p-value < 0.05 indicated by “*”

We further examined to see whether for individual genes impacted by abnormal ME-transcripts the rate of abnormal splicing form is differentially expressed among the sample groups. As shown in Fig. 3F, while the ME-transcripts of *SCN1A* gene were observed in all sample groups, its rate in percentage of the total transcripts was shown to be more than 2 times higher in the TLE-HS and TLE groups than the control group despite not having statistical significance, possibly due to the small sample size for the TLE and TLE-HS groups and/or the large relatively variations within the groups. Notably, the rate of the ME-form in the TLE-NC was shown to be statistically lower than all other 3 groups (Fig. 3F). Interestingly also is that the TLE and TLE-HS groups showed a significantly lower expression level of the normal *SCN1A* form than the control and TLE-NC group (Fig. 3G). This is also true for the total level of expression (Fig. 3H), likely due to the high ratio of these abnormal splice variants, which are likely targeted by NMD. Due to the presence of NMD, we could suspect that the observed rate of the ME-form might be under-representing what actually happened, and this could explain the overall lower expression of the genes in the epileptic groups.

## 4. Discussions

The analysis of molecular pathways contributing to epilepsy has been quite challenging due to its extremely high level of clinical and genetic heterogeneity. To date, established contributing factors include brain injury, tumor, infection, and genetic factors ^32^. For genetic factors, germline mutations of genes related to a variety of functions ranging from ion channels, enzymes, transporters, and membrane trafficking, etc., have been associated with epilepsy ^5^. Limited analysis of whole genome and transcriptome analyses have been performed for epilepsy using bulk brain tissues or laser micro-dissected tissues of specific brain structures/cell types ^28, 29, 33^.

Aside from germline mutations, a limited number of other factors, including ncRNA (e.g., miRNA)^34, 35^, somatic L1 transposition ^27^, and free L1 RNA and cDNA have been presented as possible mechanisms in contributing to neural diseases if not specifically for epilepsy ^24, 36^. For example, increased levels of L1 transcripts have been observed in many types of degenerative neuronal diseases, including frontotemporal lobar degeneration (FTLD) and Alzheimer (reviewed by Suarez et al.^37^).

In this study, we aimed to examine the role of mobile elements in epilepsy at the transcriptome level, and we focused on datasets originated from microdissection to focus on the analysis of dentate gyrus in the hippocampal tissues of the brain for TLE with and without sclerosis ^28^, using similar samples from healthy individuals as controls. The dentate gyrus is known to play a critical role in learning and memory as the primary site of adult neurogenesis in many species ^38^, and it is found to be most often involved in hippocampal neuronal loss in TLE ^39^. We argue that by focusing on this specific brain structure and cell type important for epilepsy by utilizing the laser micro-dissected samples rather than regional bulk tissue samples, we might be able to better observe patterns relevant to the disease. Nevertheless, we did also include a bulk tissue dataset for the neocortex of TLE patients ^29^ as a control for a brain region that is not related to epilepsy. We did not include the datasets for the matching hippocampal tissues of this particular study for two reasons: the epilepsy patients were not subgrouped into TLE with and without sclerosis and the samples were not micro-dissected but bulk tissues.

Among genes reported to be associated with epilepsy, the *SCN1A* gene, which encodes type 1 sodium channel alpha subunit, stands out as for being most highly associated with epilepsy ^40^. Its mutations are associated with a spectrum of phenotypes ranging from the mild form of generalized epilepsies ^41, 42^ to the extremely severe form of the Dravet syndrome ^43, 44^. Mutations in this gene is responsible for more than 60% of Dravet syndrome ^45^. In a recent study, SCN1A was shown to have lower expression in TLE-HS patient and the controls ^33^. In our study, *SCN1A* was shown to be involved in ME-transcripts with a higher rate of the abnormal splice form in the TLE and TLE-HS group with the retaining ME interrupting the open reading frame with a premature stop codon. This leads to a severely truncated protein product, missing more than 1000 amino acid residues in the C-terminus, which contains three functional domains including the sodium ion transport-associated domain and two Ion transport protein domains (Fig. 3E). The other possible outcome of this abnormal splicing is the degradation of the ME-transcripts by NMD. In both scenarios, the end result would be a loss of function for the gene or the lower expression of the gene since the ME-transcript form counted only part of all transcripts for the gene. In this case, even though there is no sequence mutation observed, the level of functional gene product would be reduced.

Regulation of alternative splicing involving two in-frame mutually exclusive exons in *SCN1A* (exon 5A and 5N) and its paralogs has been reported to be contributing to the different activities of these Na channels, which in turn impact the threshold and duration of seizure ^46, 47^. Genes known to be involved in regulating alternative splicing of these Na channel genes include NOVA in mammals and pasilla in drosophila ^48, 49^, and manipulation of the alternative splicing has been proposed as an exploitable mean for providing effective seizure control ^50^. Interestingly, NOVA genes (*NOVA1* and *NOVA2*) were not seen in our list of genes involving MEs in abnormal splicing (Table S4), indicating that a different mechanism is involved in the abnormal splicing we observed.

A recent study reported differential exon usage of a list of 124 genes among different epilepsy groups, and these genes showed enrichment of biological processes including cell adhesion, immune response or response to drugs ^33^. However, in this case, the identified alternative exons represent known alternative splice variants not involving premature stop codon, notably those associated with neuroxin genes. Their gene list seems to have little overlap with our gene list shown in Table S4, and they also show enrichment of different function categories, distinct from those shown by genes associated with the abnormal splicing we observed in this study.

It is to be noted that our study is severely limited by the very small sample sizes for the epileptic groups for being 14 and 8 samples for only 7 and 5 subjects, respectively. On one hand, being able to observe the pattern with statistical significance with such small sample sizes suggests the signal is very strong and much stronger may be observed with large sample sizes. On the other hand, it is also possible that pattern could represent something unique to this set of epilepsy samples. Nevertheless, if it can be further confirmed with additional sample sets, it not only can shed new lights on the molecular mechanism underlying epileptogenesis, but may also provide new means of epilepsy diagnosis/prognosis and treatment. It is also possible that abnormal splicing may serve as a general mechanism leading to loss of function in genetic diseases. In addition to validation studies involving large samples, future studies may extend to larger scale studies covering not only large numbers of samples but also more epilepsy subtypes to confirm the pattern we observed in this study. More in-depth and/or targeted gene studies confirming the functional impact on the impacted genes, in particular those with known association to epilepsy or neural functions, will be highly desirable. Certainly, it would be very interesting to identify the upstream contributing factors and mechanisms leading to the higher rate of ME-associated abnormal splicing.

## Supporting information

SummentalMethods_Results

## Data availability

All raw RNA-seq datasets used in this study are made available in NCBI SRA database by their respective original owners as listed in Table 1 and S1. The Perl script used to identify ME-transcripts based on the RepeatMasker annotation of the assembled transcripts, the blat output of aligning the transcripts to the reference genome and the genome annotation is available at GitHub at https://github.com/pliang64?tab=repositories.

## Supplementary data

Additional information on methods and data described in this article is provided in the supplemental file.

## Author contributions

KH and PL conceived and designed the study. KH contributed to the acquisition of RNA-seq data, interpretation of data, bioinformatics and biostatistics analysis of data, and the manuscript writing. PL contributed to bioinformatics analysis and writing of in-house Perl scripts, in-depth analysis of preliminary results. All authors read, revised and approved the final manuscript.

## Author statement

None of the material presented in the article has already been published or is under consideration for publication elsewhere, including the internet.

## Declaration of Competing Interest

The authors declare no competing interests.

## Acknowledgements

This work is in part supported by grants Canadian Research Chair program, Canadian Foundation of Innovation, Canadian Natural Science and Engineering Research Council (NSERC), and Ontario Ministry of Research and Innovation to PL. Most part of the bioinformatics analysis was completed using the Compute Canada/SHARCNET high-performance computing facilities.

## A list of supplementary figures and tables

Table S1. Datasets and sample IDs used in the human epilepsy groups

Table S2. Contribution of mobile elements (MEs) to gene transcripts in human sample groups

Table S3. Contribution of mobile elements (MEs) subfamilies to gene transcripts in human epilepsy patient groups

Table S4. A list of epilepsy-associated and neural genes and those impacted by MEs in the CDS regions in the epileptic sample groups

Table S5. Frequency comparisons of epilepsy-associated and neural genes Impacted by MEs in CDS and 3′ UTR regions in human sample groups

Table S6. The expression level of transcript forms for SCN1A gene

Figure S1. Comparisons of mobile elements (MEs)’ contribution by ME class to gene transcripts in human epilepsy groups.

Figure S2. Age profile comparisons for mobile elements (MEs) in the genome and ME-transcripts in the control and TLE-HS patients.

## Notes

https://github.com/pliang64/fatools

### Competing Interest Statement

The authors have declared no competing interest.

