## Supplementary material for "Transcriptome analysis reveals higher levels of mobile element-associated abnormal gene transcripts in temporal lobe epilepsy patients": SummentalMethods_Results

**Supplemental information on Methods:**

*1. RNA-seq data and reference genome and gene annotation collection*

RNA-seq sequences for these samples were downloaded from NCBI SRA database in compressed fastq format to our local server for in-house analysis. All RNA-seq data included were generated using Illumina HiSeq 2000/2500 platforms in paired-end reads at 150 bp with the volume being around 3.0 TB for human (Total N=90) (**Table 1** and **Table S1**). The human reference genome (GRCh38/UCSChg38, Dec 2013) and the corresponding gene annotation file (refGene.txt) files were downloaded from UCSC website (http://genome.ucsc.edu).

*2. De novo transcriptome assembly, identification of ME-transcripts and MEs' gene context*

For each RNA-seq dataset listed in **Table 1** and **Table S1**, a *de novo* transcriptome assembly was performed using SOAPdenovo-Trans [1] with the default parameter settings keeping only transcripts that are 300 bp or longer.

To identify transcripts containing ME sequences, all assembled transcripts from the above step were subject to repeat annotation using RepeatMasker (version 4.1.0, http://www.repeatmasker.org) with the repeat library setting to the corresponding species. Any transcripts for known genes containing one or more MEs are referred to as ME-transcripts. The individual sequences of all ME-transcripts were extracted from the transcript assembles using the in-house tool, fatools (available at https://github.com/pliang64/fatools). Extracted ME-transcript sequences for each sample were then aligned to the corresponding reference genome sequences using pblat [2, 3] to obtain their aligned positions in the reference genomes. The blat output was processed using an in-house Perl script to generate a delimited text file, which provides the chromosomal position and classification of the MEs in the ME-transcripts and the symbols and positions of the genes in which the ME(s) reside, as well as the gene regions to which MEs correspond to. This was done based on the position of MEs in the transcripts, and the position of the transcripts in the reference genome and the location of gene exons in reference genome. For protein coding genes, the location of MEs was also categorized into subregions of the exons as 5' UTR, 3' UTR and CDS (coding sequence). The number of ME-transcripts in each sample was collected and normalized as the number of ME-transcripts per million transcripts (TPM). TPM values were also calculated for each major ME class (SINE, LINE, LTR, SVA, and DNA transposons), and for their subfamilies showing a minimum of 1 TMP with statistical analysis performed as described in section 2.5.

*3. Detailed sequence analysis of ME-transcripts*

For a selected list of epilepsy-associated genes involving MEs in CDS, we performed detailed sequence analysis of the ME-transcripts manually using the resources and utilities on UCSC genome browser (https://genome.ucsc.edu). Specifically, the entire ME-transcript sequences in fasta format were retrieved from the transcript assemblies of the corresponding samples and used to search the reference genome and locate the MEs in CDS exons as a way of validation of the predicted CDS location of the MEs and predicting the impact on protein coding. Further, for each of such ME-transcript, we extracted the 20 bp sequences flanking the junction point (10 bp on each side), at which the normal transcript and the ME transcript differentiate due to the presence of a ME for both the normal and ME-transcripts. This pair of 20 bp sequences were then used to search the entire set of RNA-seq raw reads to count the number of reads supporting each of the two transcript sequences at that specific location. These counts were collected for each sample and normalized as reads per million reads (RPM) an in-house Perl script and compared across sample groups with statistical testing done similarly as for the ME-transcript frequency.

*4. Computational analysis*

Most of the analysis for RNA-seq data was performed on Compute Canada high-performance computing facilities (https://computecanada.ca)

### Supplementary Results:

*1. Old MEs are the main players in gene transcripts*

To better understand the contributing factor(s) related to the degree of MEs' involvement in gene transcripts, we performed age profiling of these MEs based on sequence divergences from their respective consensus sequences. In this case, the higher the divergence, the older the age of the ME in the genome.

Overall, as expected, the age profile is different among ME classes with SINEs showing a youngest peak among all ME classes, followed by DNA transposons and LTRs, and then by LINEs as the oldest (**Fig. S2**). For SINEs, the age profile is pretty much identical among the three datasets, indicating their degree of involvement in ME-transcripts is simply a reflection of their age profile in the genome with no clear age biases. For DNA transposons, the two sample groups showed a younger age profile than their genome-wide profile, indicating a bias for younger elements in ME-transcripts. LTRs showed a very wide and flat peak in the genome, likely reflecting a continuous LTR domestication process and/or steady propagation during the human genome evolution. However, LTRs showed a single main peak in the control sample group somewhere in the middle for the flat peak for the genome, but two peaks in the LTE-HS group, one being younger and the other being older than the main peak in the control group (**Fig. S2**). This indicates that LTRs involved in ME-transcripts in the LTE-HS patient transcriptomes have a unique from the younger members. For LINEs, the genome profile showed a clear peak towards the old age and the control group showed two peaks both younger than that of the genome profile, while the TLE-HS group showed a main peak aligned to the older peak in the normal group and a secondary peak at a much younger age, which happens to be the same as age of the SINE peak. This indicates that LINEs in ME-transcripts in the TLE-HS group also have a proportion contributed from young members (**Fig. S2**). Overall, the fact that no SVA was observed in ME-transcripts agrees with the general pattern observed in other ME classes.

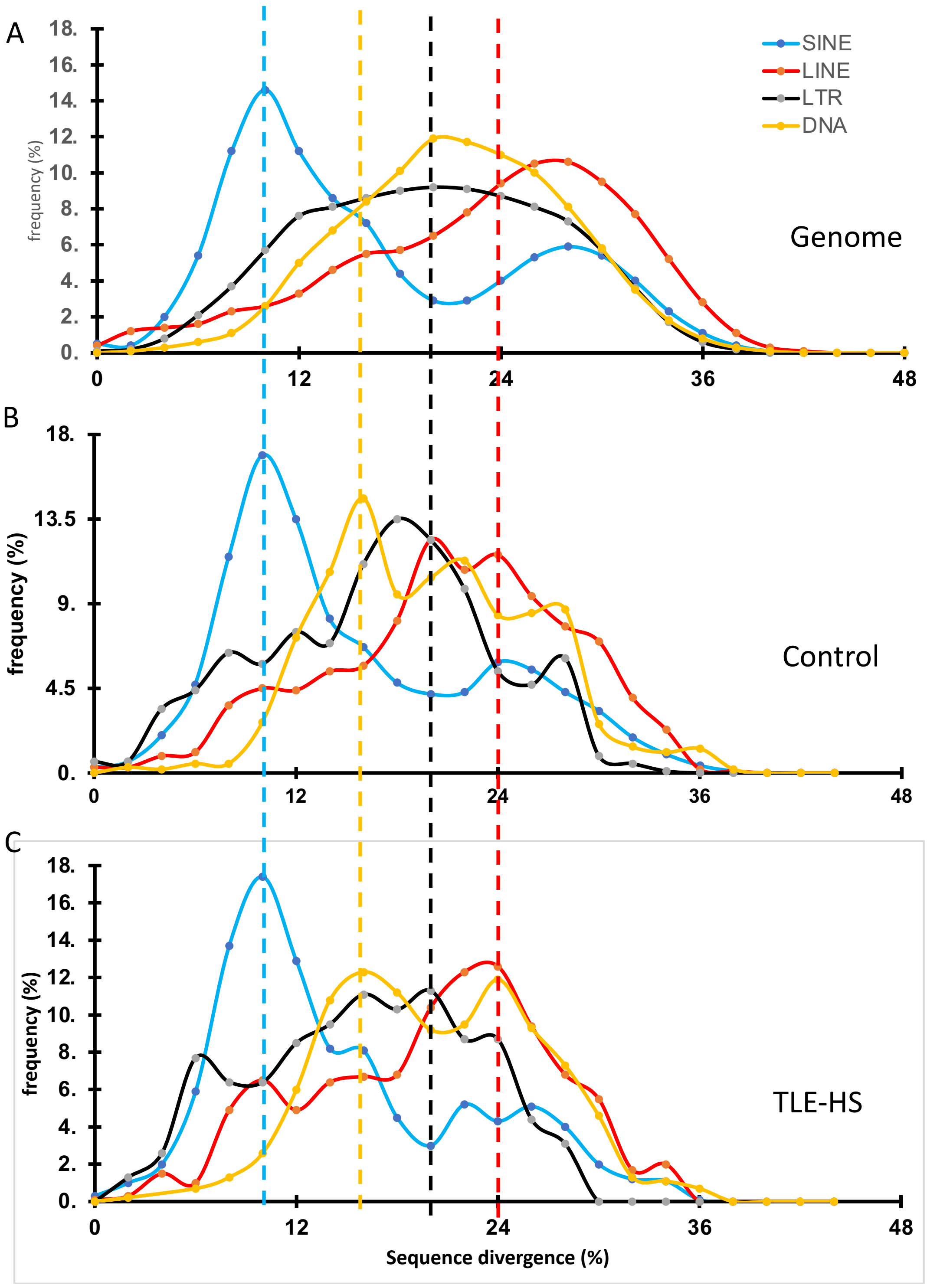
A
B
C
SINE
LINE
LTR
DNA
Genome
Control
TLE-HS
frequency (%)
Sequence divergence (%)
0
12
24
36
48
18.
16.
14.
12.
10.
8.
6.
4.
2.
0.
13.5
9.
4.5

**Figure S2. Age profile comparisons for mobile elements (MEs) in the genome and ME-transcripts in the control and TLE-HS patients.**

Line plots showing the age profiles of MEs in different class based on the percentage of sequence divergence from the corresponding ME consensus sequences as reported by RepeatMasker, with lower divergence indicating younger age. A. All MEs in the human reference genome (NCBI GRCh38); B, MEs in ME-transcripts from the control sample group; C. MEs in ME-transcripts from TLE-HS sample group.  Colour legend for B and C is the same in A. Dashed vertical line indicates the peak of the age distribution for the corresponding ME class based on colour in TLE-HS.
