## Supplementary figures and images for "Transcriptome analysis reveals higher levels of mobile element-associated abnormal gene transcripts in temporal lobe epilepsy patients"

### SupplementalFigures

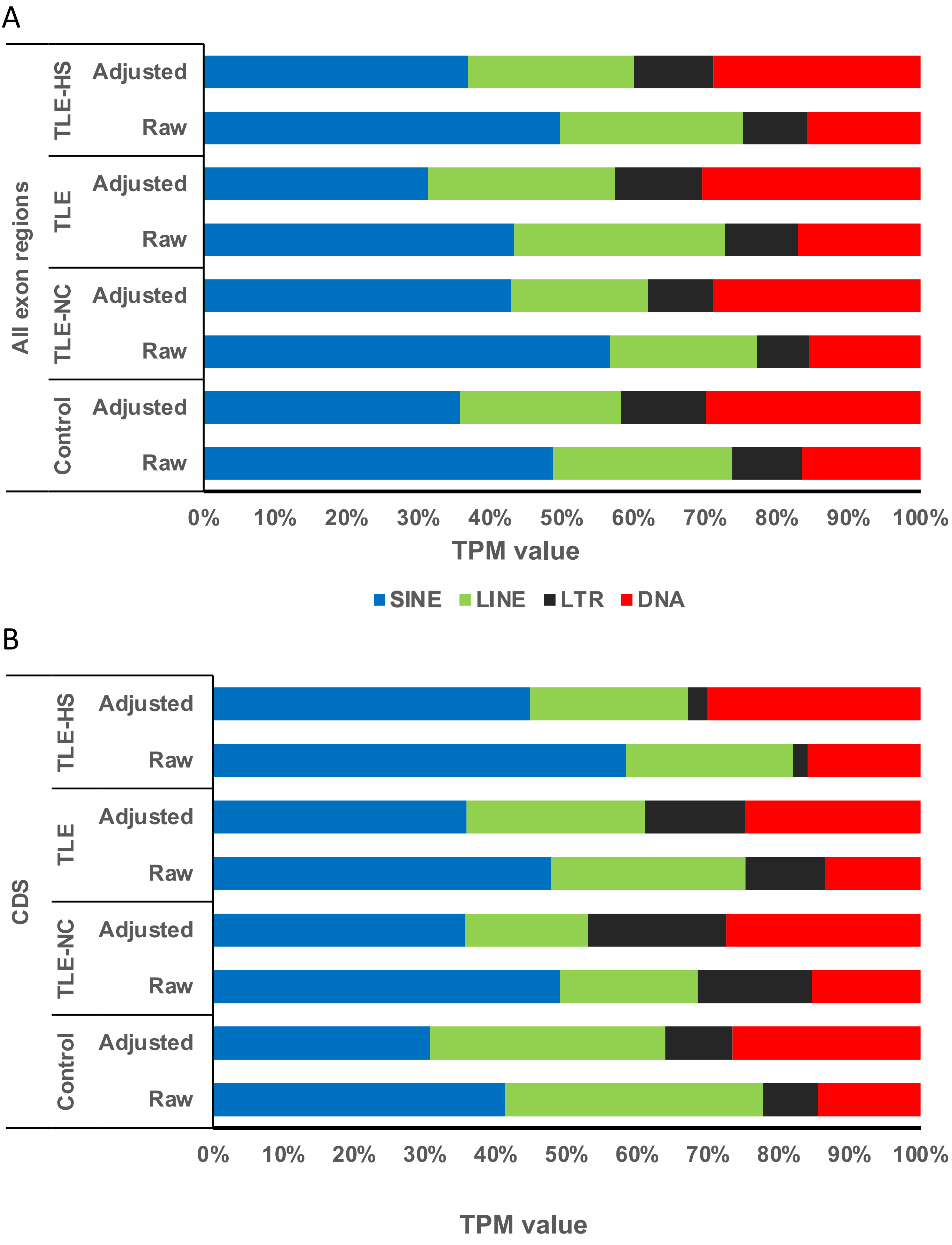

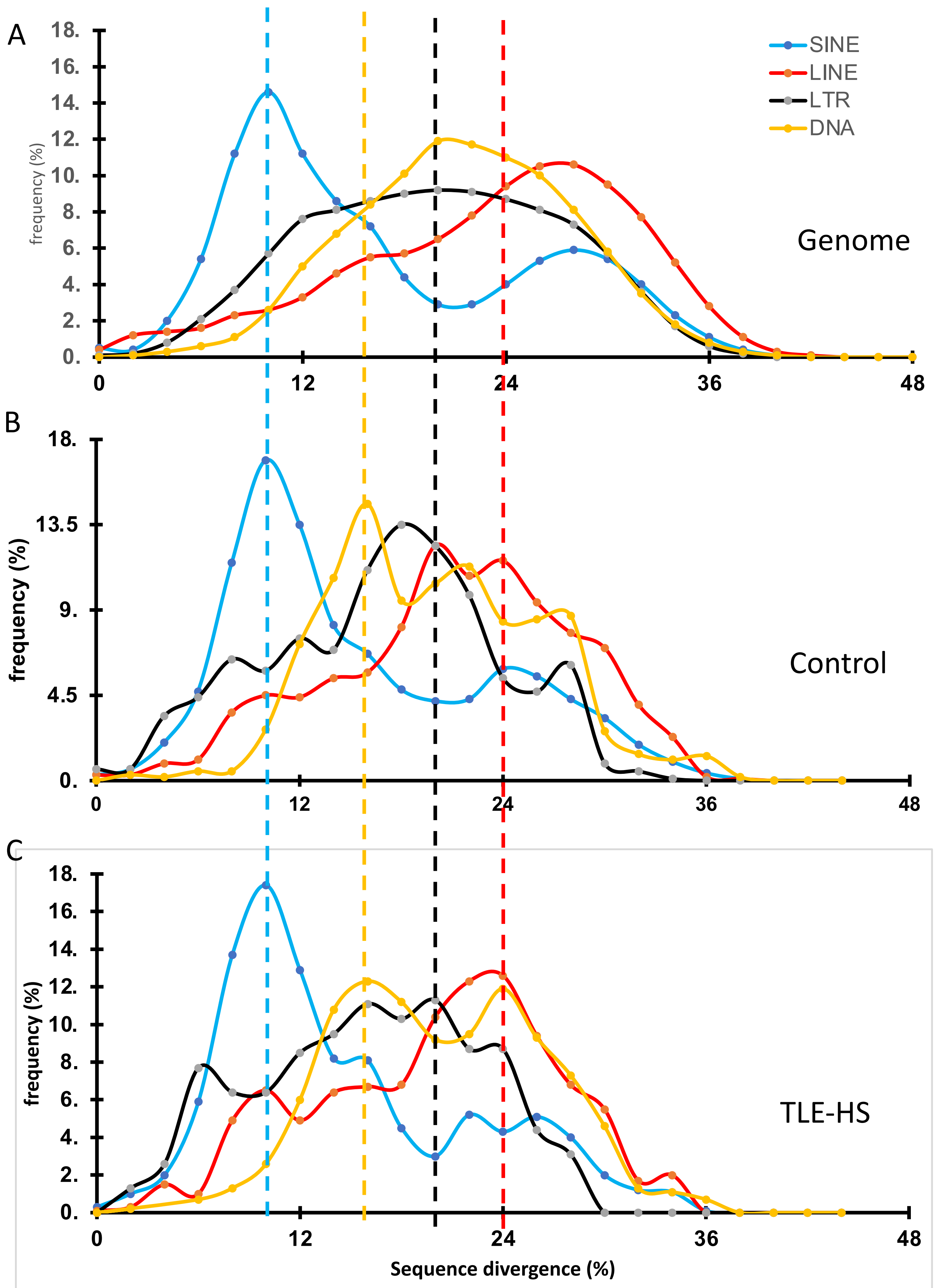
